# Ploidy tug-of-war: evolutionary and genetic environments influence the rate of ploidy drive in a human fungal pathogen

**DOI:** 10.1101/084467

**Authors:** Aleeza C. Gerstein, Heekyung Lim, Judith Berman, Meleah A. Hickman

**Affiliations:** Department of Genetics, Cell Biology & Development, College of Biological Sciences, University of Minnesota, USA; Department of Microbiology & Immunology, Medical School, University of Minnesota, USA; Department of Molecular Microbiology and Biotechnology, George S. Wise Faculty of Life Sciences, Tel Aviv University, Israel; Emory University, Department of Biology, O. Wayne Rollins Research Center, 1510 Clifton Road NE, Atlanta, GA 30322

## Abstract

Variation in baseline ploidy is seen throughout the tree of life, yet the factors that determine why one ploidy level is selected over another remain poorly understood. Experimental evolution studies using asexual fungal microbes with manipulated ploidy levels intriguingly reveals a propensity to return to the historical baseline ploidy, a phenomenon that we term ‘ploidy drive’. We evolved haploid, diploid, and polyploid strains of the human fungal pathogen *Candida albicans* under three different nutrient limitation environments to test whether these conditions, hypothesized to select for low ploidy levels, could counteract ploidy drive. Strains generally maintained or acquired smaller genome sizes in minimal medium and under phosphorus depletion compared to in a complete medium, while mostly maintained or acquired increased genome sizes under nitrogen depletion. Surprisingly, improvements in fitness often ran counter to changes in total nuclear genome size; in a number of scenarios lines that maintained their original genome size often increased in fitness more than lines that converged towards diploidy. Combined, this work demonstrates a role for both the environment and genotype in determination of the rate of ploidy drive, and highlights questions that remain about the force(s) that cause genome size variation.

## Introduction

Baseline ploidy levels vary among closely related extant species and whole genome sequencing has revealed widespread paleopolyploidy throughout the eukaryotic tree of life (reviews on animals: Gregory TR and Mable 2008; fungi: Albertin and Marullo 2012; and plants: Wendel 2015). Ploidy variation has the potential to affect many aspects of evolutionary dynamics and there are theoretical advantages to both high and low ploidy levels, primarily related to differences in the mutation rate and the efficiency of selection (Thompson and Lumaret 1992; Orr and Otto 1994; Otto and Whitton 2000; Gerstein and Otto 2009, and references within). In brief, more mutations arise in higher ploidy backgrounds, thus if new beneficial mutations are the rate-limiting step in adaptation, higher ploidy populations should be advantageous. However, if the adaptive mutations are not completely dominant or have low penetrance, they will take longer to reach high frequency in higher ploidy populations and may be lost due to chance. Additional complexities regarding the influence of ploidy level on the characteristics of beneficial mutations have been revealed in recent experimental studies with yeast. Different ploidy levels may have different suites of mutations available to them (Orr and Otto 1994; Gerstein and Otto 2009; Selmecki et al. 2015). Furthermore, the same mutations can have different effect sizes in different ploidy backgrounds, independent of dominance (Gerstein 2012; Selmecki et al. 2015), implying that ploidy itself has important consequences on the mutational pathways available to evolution.

Ploidy also has the potential to directly or indirectly influence the physiological characteristics of an organism. In single-celled eukaryotes, ploidy directly influences the volume of the cell (likely mediated through differences in nuclear volume, Schmoller and Skotheim 2015), and ploidy/cell volume have been directly implicated in differences in amino-acid pools and enzyme activity (Weiss et al. 1975), gene expression (Galitski et al. 1999; Storchová et al. 2006; Li et al. 2010; Wu et al. 2010), and protein levels (Godoy et al. 2008; Gómez et al. 2009). Polyploidy is also linked to similar phenotypic effects in multicellular organisms. Polyploid plants and animals tend to have larger cells than diploids (review of plants: Ramsey & Ramsey 2014, review of animals: Neiman et al. 2017). Compared to diploids, many polyploid plants also are larger overall, have sturdier leaves and reproductive structures, and flower later (Ramsey & Ramsey 2014). Polyploid animals have been less well characterized than polyploid plants, though also tend to be larger (Neiman et al. 2017). In all kingdoms, however, it is clear that the link between ploidy and physiological characteristics (and hence ploidy and fitness) is complex, and although generalizations can be made, some taxa do not follow the general pattern (Soltis et al. 2016, Neiman et al. 2017)., Thus, the theoretical “optimal” ploidy level may depend heavily on the specific organism (and its genetic background) and the selective environment.

Experimental evolution studies *in vitro* using ploidy variants of fungal microbes reveals an intriguing ploidy phenomenon: convergence towards the species historical, baseline ploidy within tens to hundreds of generations under diverse environmental conditions. This phenomenon has been repeatedly observed across evolutionarily divergent fungal taxa in experiments that track ploidy. These studies include ascomycete and basidiomycete fungal species whose historical baseline ploidy is diploid, including the standard laboratory yeast *Saccharomyces cerevisiae* (Gerstein et al. 2006; Gresham et al. 2008; Voordeckers et al. 2015) and human pathogenic *Candida* species (*Candida albicans*: Hickman et al. 2015; *Candida tropicalis*: Seervai et al. 2013), and in species whose historical baseline ploidy is haploid (*Cryptococcus neoformans*: Gerstein et al. 2015; *Schizosaccharomyces pombe*: V. Perrot et al., pers comm; *Aspegillus nidulans*: Schoustra et al. 2007).

That organisms typically revert back to their baseline ploidy is, at first glance, perhaps unsurprising. Eukaryotic single-celled microbes are known to have incredibly labile genomes that gain and lose chromosomes rapidly under certain environmental conditions (Gerstein and Berman 2015, and references within). However, the studies documenting ploidy convergence back to baseline raise many questions about the cellular mechanisms enabling ploidy variants to arise outside of the sexual cycle. Ploidy doubling can occur through whole genome duplication via endoreplication, which results from mitotic defects when a cell undergoes DNA replication yet fails to segregate during cell division (Otto and Whitton 2000). A mechanism for mitotic ploidy reduction (i.e. the ‘parasexual cycle’, Pontecorvo 1956; Bennett et al. 2003) remains mostly uncharacterized, particularly when the rate of chromosome loss is too rapid for selection to be implicated (i.e., when an entire set of chromosomes is lost in a very small number of generations, e.g., Gerstein et al 2008). Furthermore, in the myriad of experiments that have observed convergence to baseline ploidy, a clear advantage attributable to ploidy change *per se* has only been demonstrated once (Venkataram et al. 2016, in *S. cerevisiae* adapting to glucose limitation). It thus remains elusive what evolutionary mechanisms enable ploidy variant cells to sweep through a population of non-baseline ploidy cells. Hence, we term the force acting on a lineage to asexually revert back to its baseline ploidy level as ‘ploidy drive.’

Ploidy diversity among isolates has previously been observed in both environmental and clinical populations of *S. cerevisiae* (Ezov *et al*. 2006; Zhu, Sherlock, Petrov 2016). This type of ploidy variation is also observed in the opportunistic human pathogen *C. albicans* (Suzuki et al. 1982; Marr et al. 1997; Hickman et al. 2013; Abbey et al. 2014), a budding yeast that diverged from *S. cerevisiae* prior to the whole-genome duplication (Diezmann et al. 2004; Kellis, Birren and Lander 2004). The isolation of natural ploidy variants suggests these atypical ploidy individuals may play an important evolutionary role. An evolutionary link between ploidy and ecological environment has previously been hypothesized in the context of nutrient limitation (Lewis 1985; Elser et al. 1996; Sterner et al. 2002; Elser et al. 2007). The nutrient limitation hypothesis suggests that as haploid cells have a higher surface area: volume ratio (SA: V) relative to higher ploidy cells, nutrient scarcity serves as a selective force promoting haploidy because of the increased capacity for passive nutrient uptake from the extracellular environment. An increase in energy and material costs is also predicted to select against higher ploidy under nutrient scarcity, particularly when there is selective pressure on growth rate due to the high demand for phosphorus for ribosomes and nitrogen for proteins (Hessen et al. 2010).

Testing the nutrient limitation hypothesis in fungal taxa has been limited to *Saccharomyces*, with extremely mixed results (Adams and Hansche 1974; Glazunov et al. 1989; Mable 2001). Recently, a comprehensive study examined 24 *S. cerevisiae* and 27 *S. paradoxus* isogenic haploid and diploid natural isolates in 33 distinct environments and found that although environment × ploidy interactions were frequent, there was no clear-cut benefits associated with haploidy under nutrient limitation (Zörgö et al. 2013). As a representative example, diploidy was advantageous in two of four non-preferred nitrogen sources, with no difference between haploids and diploids in the other two environments. What has not been tested, however, is whether ploidy drive can be mitigated by nutrient limitation.

We tested the influence of nutrient limitation on ploidy drive by evolving clinical and laboratory haploid, diploid, and polyploid strains of *C. albicans* in complete medium and under nutrient limitation. *C. albicans* provides a compelling study system because the primary site of colonization is as a commensal organism in the human gut (Odds 1988), where it experiences general nutrient limitation through competition with other microbial organisms. Furthermore, *C. albicans* proliferates in a large diversity of sites with the human body, including kidneys, skin, nails, and mucosal surfaces (Odds 1988). In the laboratory, a variety of stress conditions result in karyotypic changes (Janbon et al. 1998; Bennett and Johnson 2003; Wellington and Rustchenko 2005; Chang et al. 2014; Harrison et al. 2014), suggesting that a labile genome is one way *C. albicans* deals with the large breadth of ecological niches and environmental challenges it commonly faces. If the nutrient limitation hypothesis has a significant effect on ploidy drive, we predicted that compared to a complete medium, under nutrient limitation initially haploid strains should stay haploid, while initially polyploid strains should converge to diploidy faster. Unexpectedly, we found that a more nuanced view of the influence of nutrient limitation on ploidy drive was required. Strains evolved under phosphorus depletion and minimal medium generally behaved as predicted, while strains evolved under nitrogen depletion behaved more similarly to those evolved under complete medium.

## Methods

### Strains & environments

We utilized nine *C. albicans* strains for this study that vary in their relationship to each other (Table 1). The first set are homozygous lab strains, consisting of two related haploid strains (1N1 & 1N2, that are ~91% similar) and a diploid strain (2N1) that is isogenic to strain 1N2. The second set of strains is of clinical origin, and contains a heterozygous diploid strain (2N2) and a polyploid strain with a complex karyotype (4N1) isolated from the same patient, and a euploid tetraploid strain isolated from a different patient (4N2). The remaining strains include the diploid laboratory reference strain SC5314 (2N3), and two related polyploids (4N3 & 4N4). All strains have previously been genotyped by either CGH-SNP arrays or whole genome sequencing (see Table 1 for references).

**Table 1.**
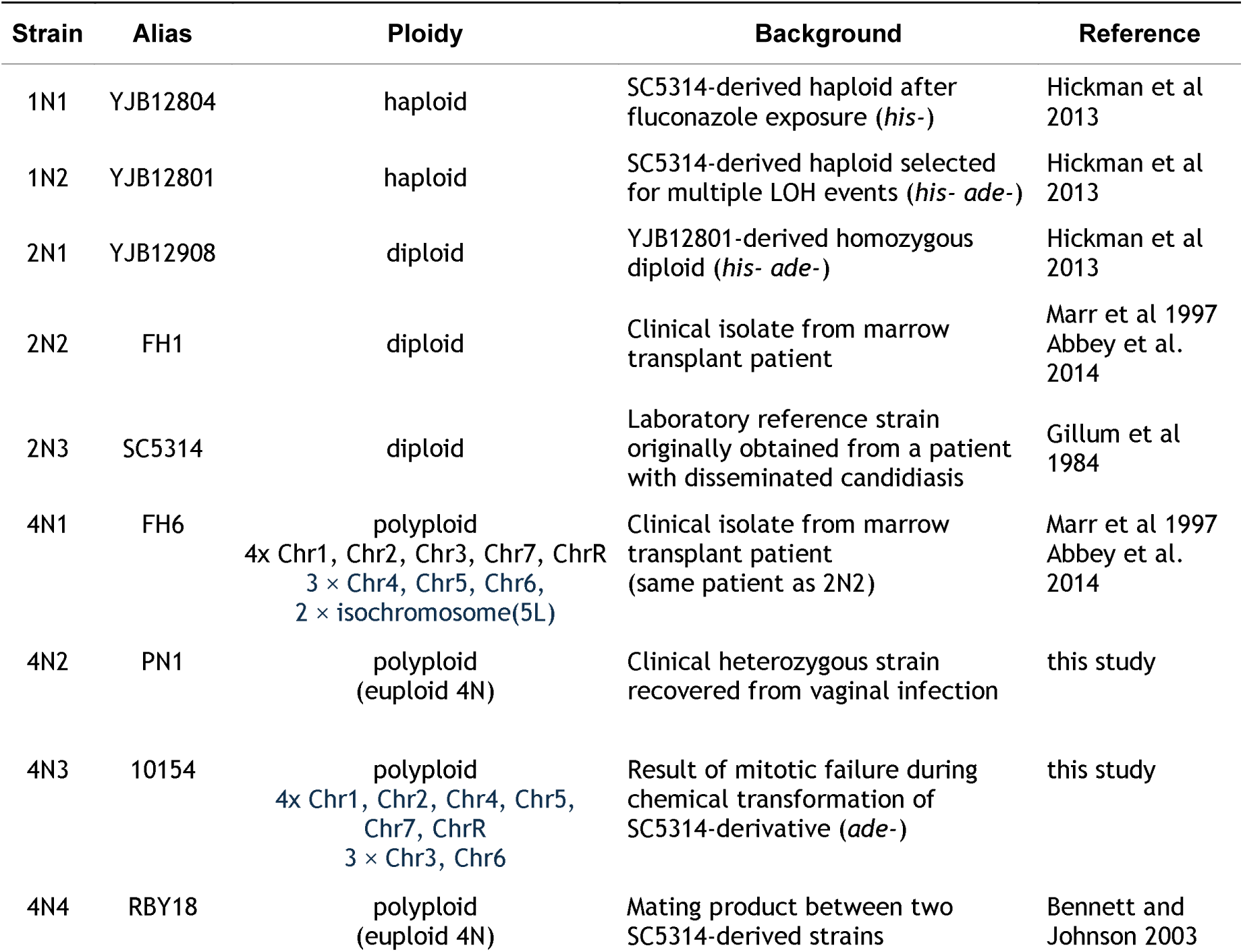
Strains used in this study. Genomic structure of previously unpublished strains determined from SNP-CGH array (4N2; see Hickman et al. 2013 for methods) and Illumina whole-genome sequencing (4N3; see Abbey et al. 2014 for methods).

Each strain was initially streaked from glycerol stocks stored at −80°C onto an SDC (synthetic defined complete medium) plate. After 48 h at 30°C, a single colony was arbitrarily chosen and transferred to a microcentrifuge tube in 15% glycerol. We refer to these single-genotype stocks as the “ancestral strains.”

We assessed ploidy drive in four different environments. Standard complete yeast medium (SDC) contains 1.7 g/L yeast nitrogen base (which includes the phosphorus source, 1.0 g/L KH_2_PO_4_), 5.0 g/L ammonium sulfate (the primary nitrogen source), 0.08 g/L uridine, 0.04 g/L adenine, and standard amino acids (Rose et al. 1990). The three nutrient reduction environments were MM (complete medium without amino acids, i.e., a general minimal medium), phosphorus-depleted (Pdep) and nitrogen-depleted (Ndep). 2% dextrose was added to all environments as the primary carbon source, the standard amount used for optimal yeast growth (Amberg et al. 2005).

Preliminary experiments were conducted to determine the maximal amount of nitrogen and phosphorus that could be removed while still achieving a growth plateau by 48 hours (results not shown). Ndep was constructed with the same base medium as MM without adding any ammonium sulfate (i.e., 100% reduction of the primary nitrogen source). For Pdep we found that we could not completely eliminate the primary phosphorus source (KH_2_PO_4_) as strains begun to go extinct after three transfers. We thus constructed Pdep medium with YNB without KH_2_PO_4_ (purchased from Sunrise Scientific Products) and added back 10% of the typical amount of KH_2_PO_4_ (0.1 g/L; i.e. 90% reduction of the phosphorus source). We also added 0.493g/L KCl to Pdep to compensate for the potassium lost from reduction of KH_2_PO_4_ (following Homann et al. 2009). Due to auxotrophic strains in the strain set (Table 1), we were required to supplement MM, Pdep, and Ndep with 20mg/L adenine and 20mg/L histidine, thereby adding a secondary, albeit minor, nitrogen source beyond ammonium sulfate.

### Batch culture evolution

We conducted batch culture evolution experiments in SDC, MM, Pdep and Ndep in three batches (due to incubator capacity): 1) SDC, 2) Pdep, and 3) MM and Ndep. To initiate each evolution experiment batch, the nine ancestral strains were streaked from frozen glycerol stocks onto SDC plates. After 24 h at 30°C, a single colony from each ancestral strain was transferred to 4 mL SDC and incubated shaking at 250 rpm at 30°C for 24 h. Overnight culture OD was standardized to the lowest OD, and 10 µL was inoculated into 1 mL of the appropriate medium in deep well culture blocks. For each strain we inoculated twelve wells, yielding twelve independent evolutionary lines per strain and environment.

We passaged all lines for 21 transfers in each environment with the time of transfer occurring after stationary phase was reached: for SDC, transfers were done every 24 ± 2 h; for MM, Ndep, and Pdep transfers were done every 48 ± 2 hours. For all environments we conducted 1:101 transfers of 10 µL into 1 mL of culture in deep (3 mL) 96 well culture blocks, which were incubated at 30°C with continuous shaking at 250 rpm. Ancestral and evolved samples were frozen after transfers 1 and 21 in 50% glycerol maintained at −80°C. We thus evolved a total of 432 lines (9 ancestral strains × 4 growth conditions × 12 replicate lines) for ~140 generations (21 transfers × 6.67 generations between each transfer).

### Fitness measurements of ancestral strains and evolved lines

We use term biological replicate to refer to measurements obtained from overnight culture that was grown up from different colonies, and technical replicate for culture that came from the same overnight tube but measured independently. One biological replicate was measured for growth rate of ancestral strains in all environments in the preliminary phosphorus and nitrogen experiments. The ancestral strains were streaked onto SDC plates and incubated for 48 h at 30°C. A single colony was randomly chosen and inoculated into 4 mL SDC and incubated while shaking at 250 rpm for 24 h at 30°C. Optical density (OD) at 595 nm was measured using a NanoDrop (Thermo Scientific) and sterile dH_2_O was added as necessary to standardize all overnight cultures to the same (minimum) OD. Standardized cultures were then diluted 1:101 into 1 mL of the appropriate medium. 150 µl of diluted culture was transferred to a clear round bottom 96 well plate and placed on a Tecan microplate reader, with two technical replicates per strain. OD was automatically read every 15 min for 48 h at 30°C with on the highest shaking setting.

A separate growth rate experiment was conducted on the evolved lines. All lines evolved in the same environment were prepared and measured on the same day. Due to space limitations, we measured the growth rate for only ten of the twelve evolved lines (always the first ten wells) from each strain in order to include two biological replicates of each ancestral strain. Ancestral strains were streaked onto SDC plates and incubated for 48 h at 30°C. Two colonies were randomly chosen and inoculated into 4 mL SDC and incubated while shaking at 250 rpm for 24 h at 30°C. On the same day ancestral colonies were inoculated, 10 µL of the appropriate frozen evolved culture (i.e., wells 1-10 from each line) was transferred into 500 µL SDC and incubated at 30°C for 24 h, shaking at 250 rpm. Overnight culture ODs were measured on the Tecan plate reader and standardized with sterile dH_2_O to the lowest OD. Standardized culture was diluted 1:101 into 1 mL of the evolutionary medium and two technical replicates of 150 μl were transferred to a clear round bottom 96 well plate and placed on a Tecan microplate reader. Optical density at 595 nm was recorded every 15 min for 48 h, with growth at 30°C with high shaking.

Growth rate was determined using custom R scripts that calculate the maximal growth rate in each well as the spline with the highest slope from a loess fit through log-transformed optical density data that reflects the rate of cell doubling (as in Gerstein and Otto 2011). For each ancestral strain we calculated the mean ancestral growth rate across all measured replicates for our analyses (i.e., three biological replicates, each with two technical replicates). Yield was obtained from these experiments as the optical density at 24 hours (SDC) or 48 hours (all other environments), i.e., the duration of growth between transfers.

### Genome size analysis with flow cytometry

Flow cytometry was used to determine the genome size of evolved populations from all lines. This type of flow cytometry uses a dye that stains nuclear DNA, and thus enables total nuclear genome size quantification by measurement of fluorescence intensity. Throughout, when we use the term “genome size”, we are referring to this measure, not the haploid genome size (“C-value”). Sample preparation was similar to previously published methods in *C. albicans* (Hickman et al., 2013) with the following modification: SybrGreen dye was diluted 1:100 in 50:50 TE and the final incubation step was conducted overnight in the dark at room temperature (the previously published protocol used 1:85 diluted SybrGreen solution and incubated overnight at 4°C). All ancestral and evolved lines from all environments from the same initial ploidy group were always prepared simultaneously and run on the same day on an LSRII flow cytometer (BD Biosciences); this enables us to accurately compare ancestral and evolved genome sizes. We collected FITC-A fluorescence data from 10 000 cells from each population. Cells are collected from all stages of the cell cycle, leading to some unavoidable ambiguity in ploidy determination (e.g., a haploid in the G2 phase of mitosis contains the same amount of DNA as a diploid in G1 phase). As we purposefully did not bottleneck populations prior to analysis, in some cases multiple peaks representing subpopulations of differing genome sizes were present (Figure 1). As nuclear genome size is determined by fluorescence intensity, the units are arbitrary and depend entirely on the machine settings, which were chosen to enable quantification of haploid to octoploid G1 peaks, and kept constant throughout our experiments.

**Figure 1.**
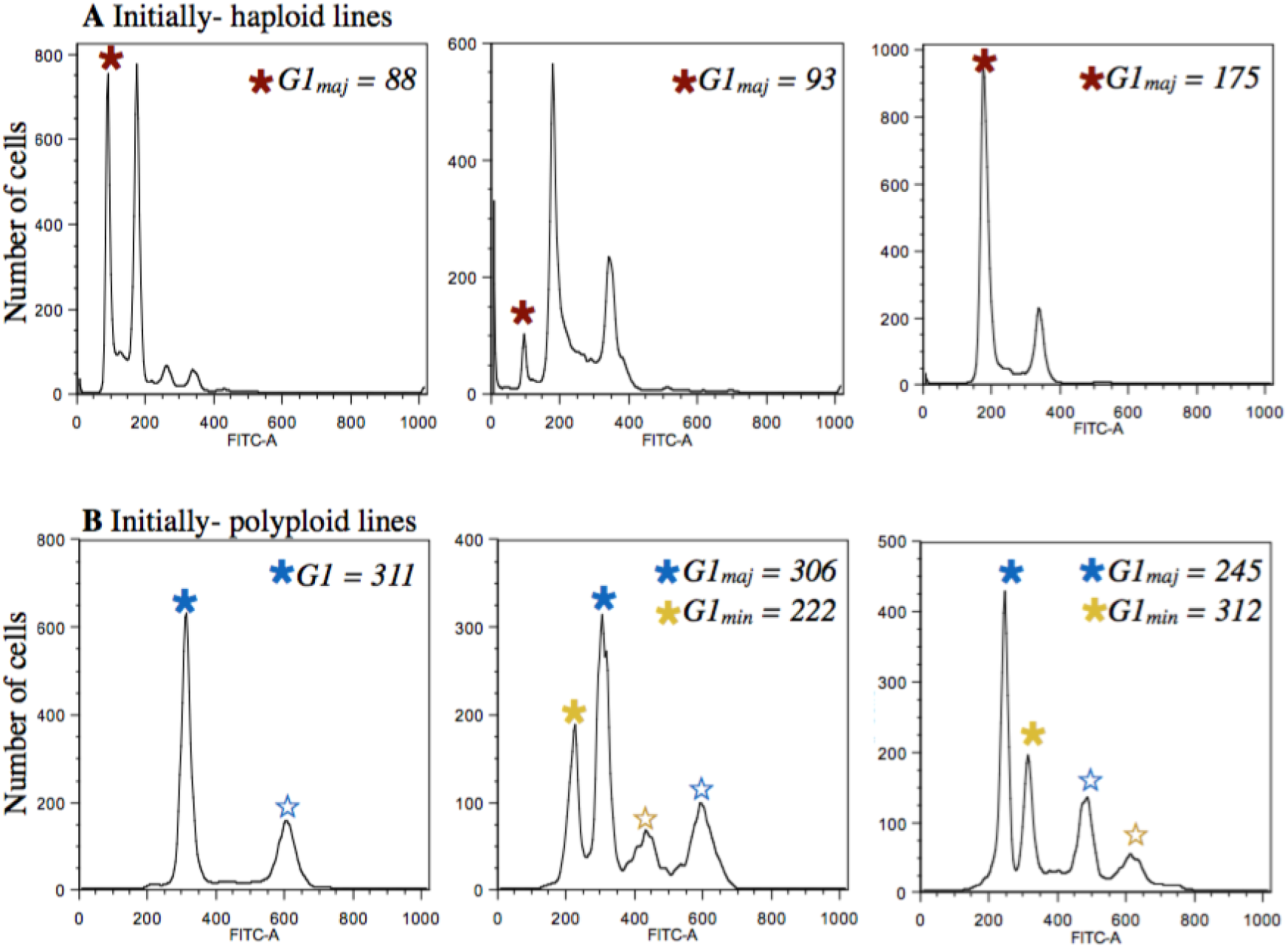
Ploidy analysis from plots of FITC intensity. A) Three representative flow profiles after evolution from initially haploid lines. In each plot the major ploidy peak (G1_maj_) is indicated by a red star. Due to ambiguity between the haploid G2 and diploid G1 peak, if a haploid G1 peak was present, regardless of height, this peak was recorded as G1_maj_ (center panel). B). Three representative flow profiles after evolution from initially polyploid lines. In each plot the mean of the major ploidy peak (G1_maj_) is indicated by a blue star. For initially diploid and polyploid lines, when more than two peaks were present (indicative of a polymorphic population), we recorded G1_maj_ from the highest peak (blue star) and G1_min_ from the next highest peak (yellow star). Note that because we assay cells at all stages of the cell cycle, we did not record the mean of peaks that are consistent with the G2 phase (open stars).

The mean G1 peak of the predominant (“major”) genome size peak was recorded for each replicate in Flowjo (Tree Star) using the cell cycle platform to fit the Watson pragmatic model. Ancestral haploid cells always contained a large G2 peak, thus when a visible haploid peak was retained after evolution we recorded the major peak for that population as haploid (Figure 1A). Conversely, ancestral diploid and polyploid cells are predominantly in G1 phase after an overnight in YPD, thus for initially diploid and initially polyploid evolved populations we recorded the major ploidy population as the highest peak (Figure 1B). All samples were prepared and measured twice by flow cytometry with extremely consistent results. We based the majority of our analyses on the major peak observed but also recorded additional (“minor”) peaks when they were present. Total nuclear genome size change was calculated by comparing the major evolved peak to the median ancestral genome size.

### Statistical analyses

All analyses were conducted in the R programming language (R Core Team 2014). Accordingly, the code used to generate all figures and conduct all analyses is available at [insert Dryad link here]. To statistically examine change in genome size as a response variable we separately analyzed strains from the three initial ploidy groups (haploid, diploid, and polyploid). The magnitude of change involved in ploidy drive from haploids or polyploids to diploidy is inherently different (one copy of the genome to two vs. four copies to two). Furthermore, as discussed in the Introduction, different mechanisms are involved in the increase or decrease of genome size, and intermediate values (aneuploidy) are only observed from initially polyploid lines.

To assess the validity of using parametric statistical tests we utilized a bootstrap technique. In each case we compared the observed test statistic to the distribution of test statistics from 1000 datasets where the response variable was randomly sampled without replacement. In all cases the position of the observed test statistic within the bootstrap distribution matched the p-value we report [link to “PloidyDrive_stat.R” file on dryad].

To examine the factors that influence genome size change we ran two-way ANOVAs with strain, evolutionary environment, and their interaction as the predictor variables and change in genome size as the response. When the interaction was not significant we re-ran the model without this term. To determine the relationship among environments we ran the HSD.test function (i.e., Tukey test with multiple comparisons) from the agricolae package (de Mendiburu 2015).

Among-strain differences for initial fitness were separately examined for each environment with an ANOVA followed by a Tukey test. To test for the correlation between initial fitness and change in fitness in each environment we used the non-parametric Kendalls’s rank association test.

To test whether initial ploidy and/or evolutionary environment significantly influenced the observed fitness changes we ran a linear mixed-effect model for each fitness measure (i.e., growth rate and yield) using the lme function from the nlme package (Pinheiro et al. 2016) with environment, ploidy and their interaction as predictor variables and strain as a random effect. The predicted marginal means, confidence intervals, and significance of contrasts between each pair of environments for each ploidy group were obtained using the lsmeans function.

Finally, to test whether change in genome size predicted improvement in fitness, for each initial ploidy group we ran a linear mixed-effect model with the change in fitness as the response variable, change in genome size as the predictor variable, and strain background and environment as random effects with the lmer function from the lme4 package (Bates et al. 2015) (degrees of freedom and p-values calculated using the Satterthwaite approximation implemented in the lmerTest package, Kuznetsova et al. 2016).

## Results

### Evolutionary environment influenced the rate of ploidy drive

After ~140 generations of evolution in synthetic defined complete medium (SDC), minimal medium (MM), phosphorus-depleted medium (Pdep) and nitrogen-depletion medium (Ndep), only ~50% (49/98) of initially haploid, and ~74% (142/192) of initially polyploid lines maintained even a minor cell population of their initial ploidy state. By contrast, 96% of diploid lines remained exclusively diploid (138/144 lines) (Figure S1). As expected, ploidy drive was thus broadly observed. To determine whether evolution under nutrient limitation specifically could counteract or enhance ploidy drive, we compared the rate of ploidy drive under the different environmental conditions.

For both diploid and polyploid strains, the change in genome size (measured as total nuclear DNA through flow cytometry) was influenced by an interaction between strain background and the environment, while in haploids only the environment was a significant predictor (Figure 2, though note that we only examined two haploid strains that are genetically related to each other; haploids—enviro: F_3,91_= 69.4, *P* < 0.0001, strain: F_1,91_= 0.02, *P* = 0.89; diploids—enviro: F_3,132_ = 7.22, *P* < 0.0001, strain: F_2,132_ = 9.16, *P* < 0.0001, strain*enviro: F_6,132_ = 4.04, *P* = 0.001; polyploids—enviro: F_3,175_ = 10.8, *P* < 0.0001, strain: F_3,1756_= 63.4, *P* < 0.001, strain*enviro: F_9,1765_= 7.7, *P* < 0.001). The precise relationship among environments depended on the initial ploidy group (Table S1). The rate of ploidy drive from haploidy to diploidy was highest in SDC, lowest in MM and Pdep, and intermediate in Ndep. Correspondingly, the rate of ploidy drive from tetraploidy towards diploidy was lowest in SDC, highest in MM and intermediate in Pdep and Ndep. Overall, strains evolved under SDC and Ndep thus tended to acquire or maintain high genome sizes, while strains evolved under MM and Pdep tended to acquire or maintain lower genome sizes. The rate of ploidy drive varied significantly among diploid strains (the only observed genome size changes occurred in strain 2N1, the homozygous diploid, during evolution in Pdep) and the polyploid strains (4N1 lines more frequently evolved towards diploidy in all environments compared to the other strains, Table S1 and Figure S1).

**Figure 2.**
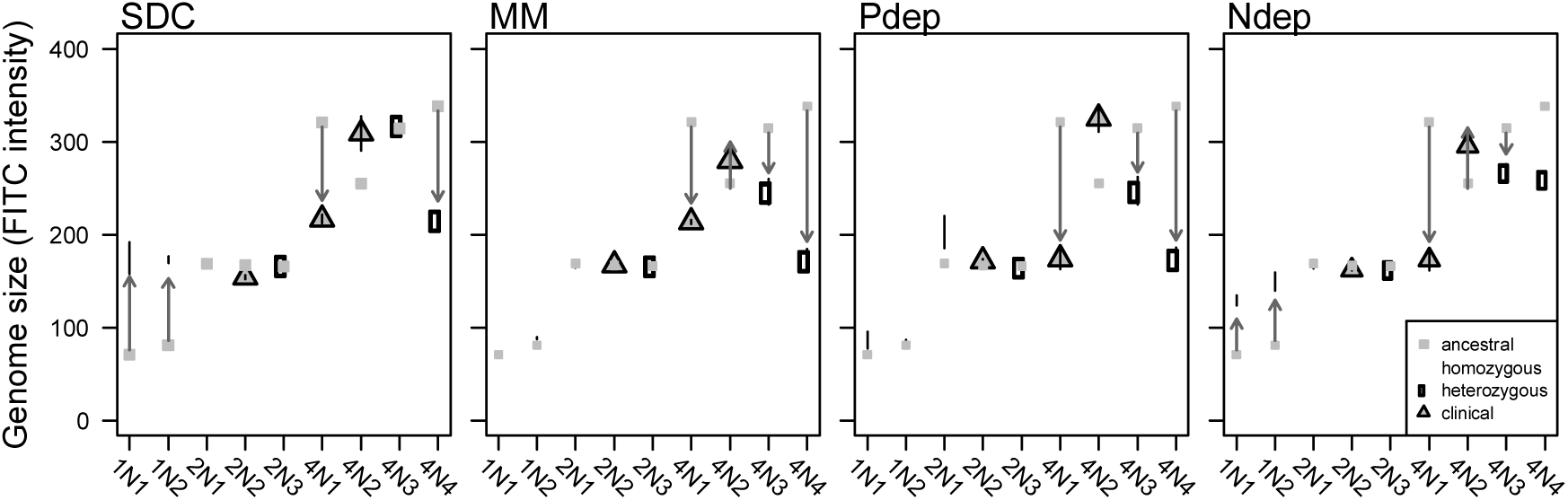
The frequency of ploidy drive was influenced by evolutionary environment and strain background. Haploid, diploid, and polyploid lines were evolved in complete (SDC), minimal (MM), phosphorus-depleted (Pdep), and nitrogen-depleted (Ndep) medium. Plotted is the mean (+/− SE) major-population genome size of 12 replicate lines evolved for ~140 generations (see Figure S1 for evolved genome size of each line individually). Genome size is measured as the fluorescence intensity in the FITC channel of SybrGreen dye. This dye stains nuclear DNA and thus fluorescence intensity reflects the total nuclear genome size (note that fluorescence intensity does not scale linearly with genome size; intensity of ~90 corresponds to haploidy, ~190 corresponds to diploidy, while ~ 310 corresponds to tetraploidy). Grey squares indicate the range of genome sizes of twelve ancestral replicates measured on the same day as evolved lines.

### Initial fitness does not predict ploidy changes

We tested whether the observed genome size changes could be explained by variation in initial growth rate or yield (optical density at the time of transfer: 24 hours in SDC, 48 hours in MM, Ndep and Pdep), two measures typically used to characterize fitness in microbes. The homozygous strains (1N1, 1N2 and 2N1) were always amongst the slowest growing and lowest yield strains for all growth environments, including MM and Pdep, the environments where haploidy was maintained (Figure 3). For the remaining heterozygous diploid and polyploid strains, strain background rather than ploidy group *per se* seemed to determine initial growth rates. Although the two heterozygous diploid strains were always among the most fit, there were no clear or consistent growth differences between them and the polyploid laboratory or clinical strains that could have predicted genome size changes. Differences in initial fitness thus did not explain the observed variation among environments for ploidy drive.

**Figure 3.**
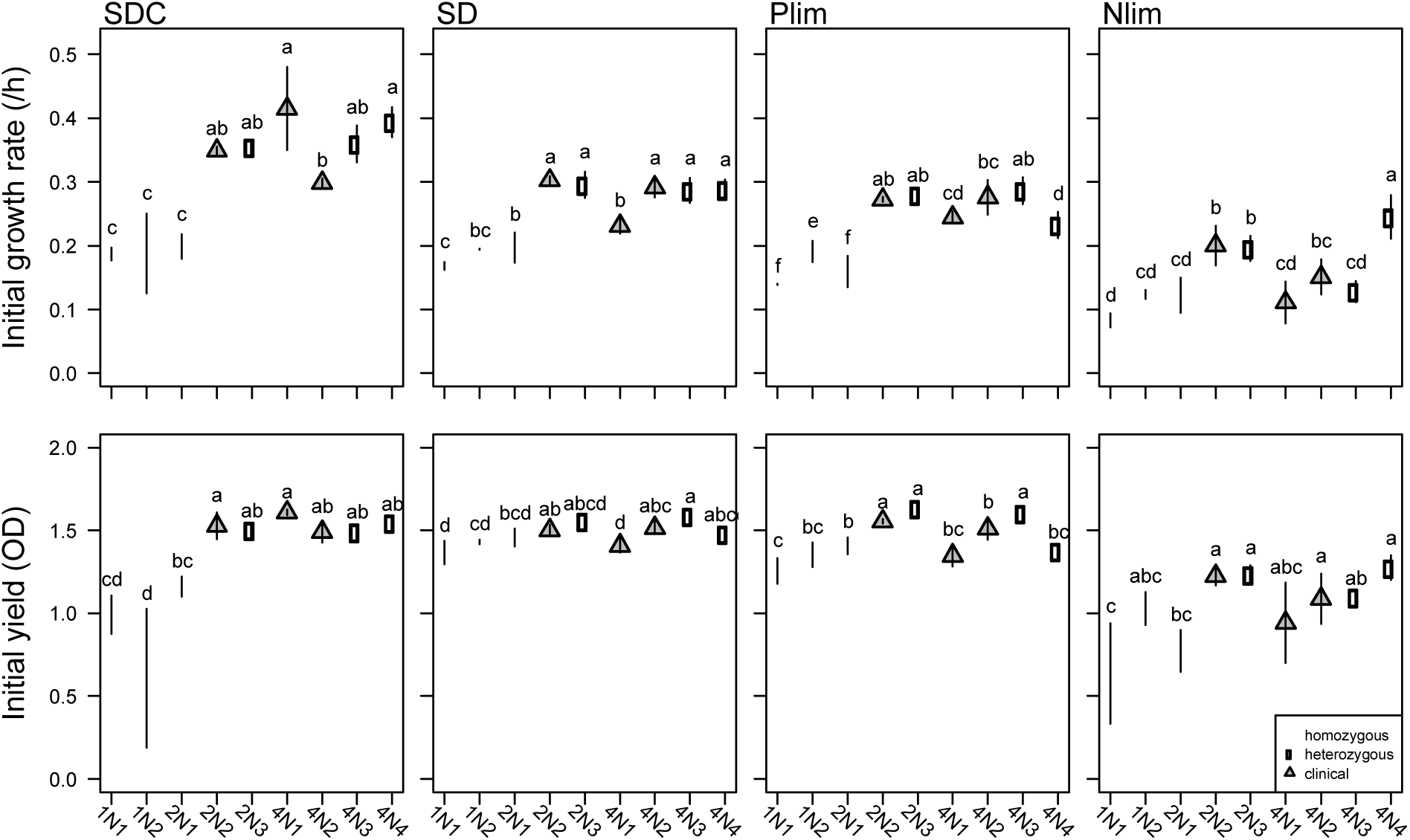
Initial ploidy does not predict initial fitness. Mean (+/− SE) initial growth rate (top row) and yield (bottom row) of ancestral strains in complete (SDC), minimal (MM), phosphorus-depletion (Pdep), and nitrogen-depletion (Ndep) medium. Yield was measured at 24 h (SDC) or 48 h (all other environments) and corresponds to the duration of growth between transfers during the evolution experiment. Letters above the symbols indicate statistical differences among-strains for each environment from a post-hoc Tukey Test; when strains do not share a letter they are significantly different from each other.

Consistent with Fisher's geometric theory of adaptation (Orr 2005), the improvement in fitness in SDC was significantly negatively correlated with initial fitness (growth rate: T = 5, *P* = 0.006, cor = −0.72; yield: T = 5, *P* = 0.006, cor = −0.72; Figure 4). A negative correlation between initial fitness and improvement in fitness was not, however, observed for either fitness measure in any other environment (growth rate—MM: T = 26, *P* = 0.12; Pdep: T = 12, *P* = 0.26; Ndep: T = 19, *P* = 0.92; yield—MM: T = 21, *P* = 0.61; Pdep: T = 15, *P* = 0.61; Ndep: T = 13, *P* = 0.36).

**Figure 4.**
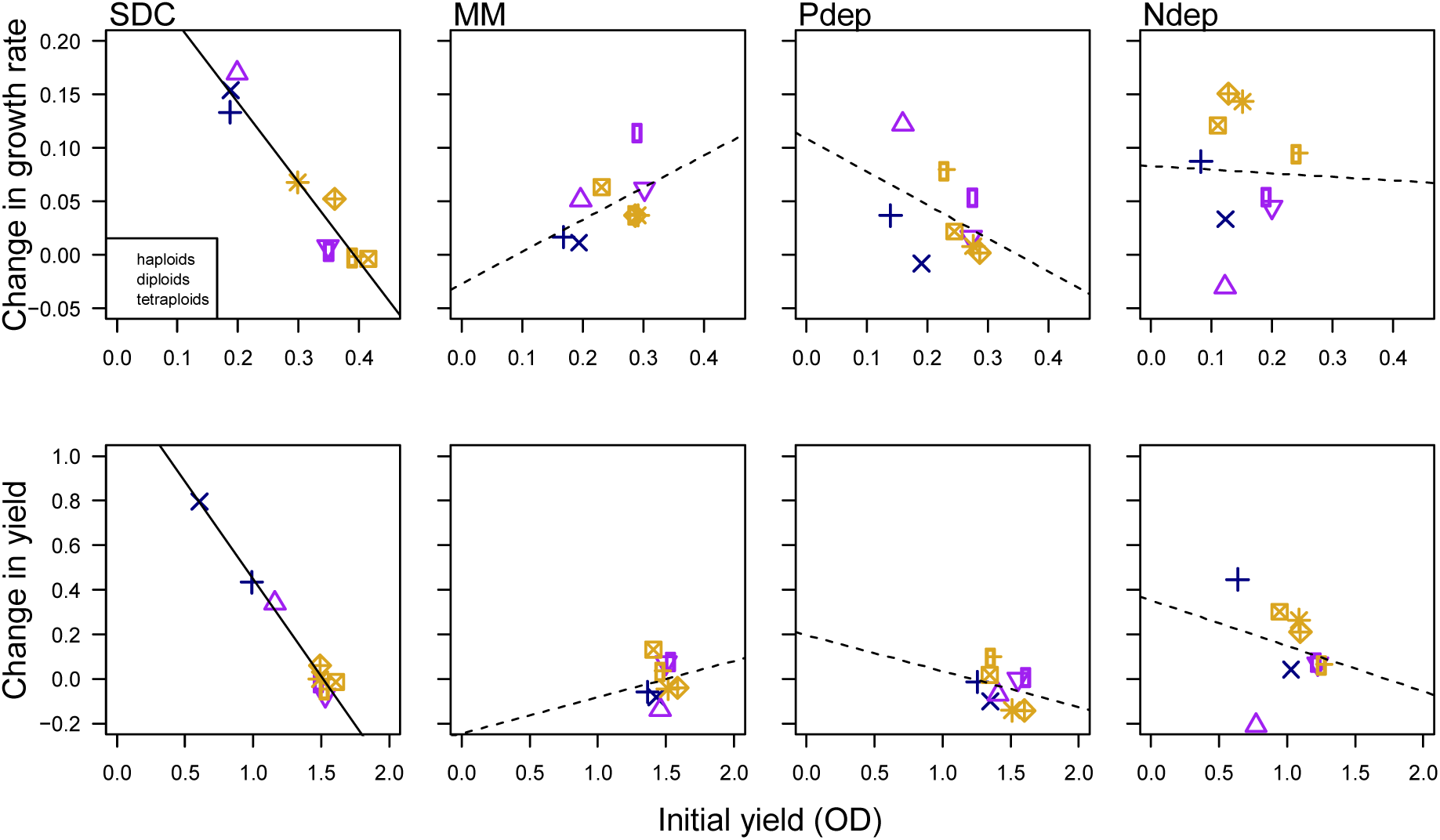
Changes in fitness do not generally correlate with initial fitness. Initial fitness measured as growth rate (top) and yield (bottom) is negatively correlated with change in fitness in SDC; the other environments show no significant correlation. A solid line indicates a significant (p < 0.05) correlation.

### Changes in genome size and improvement in fitness are not correlated

The majority of evolved lines increased in growth rate and yield relative to the ancestral strains (Figure S2). Across the entire dataset, the fitness changes were significantly influenced by an interaction between initial ploidy and evolutionary environment (growth rate—ploidy: F_2, 6_ = 0.01, *P* = 0.99; environment: F_3, 342_ = 14.51, *P* < 0.0001; ploidy*environment, F_6, 342_ = 37.4, *P* < 0.0001; yield—ploidy: F_2, 6_ = 14.19, *P* = 0.005, environment: F_3, 342_ = 39.84, *P* < 0.0001, ploidy*environment, F_6, 342_ = 36.6, *P* < 0.0001).

The mean fitness improvement compared among environments differed for the initial ploidy groups (Figure 5, statistical results in Tables S2 & S3). The growth rate improvement in haploid strains mirrored that of genome size change: they improved the most in SDC, the least in MM and Pdep, and an intermediate amount in Ndep. The growth rate of diploid strains increased significantly less in Ndep compared to other environments. In sharp contrast, polyploid strains improved growth rate significantly more in Ndep (the environment where genome size changed the least), compared to all other environments. Improvements in yield production had the same pattern as growth rate improvement in haploids and polyploids. Diploid lines showed very little change in yield, with lines evolved in Pdep and Ndep showing a significant reduction in yield compared to SDC.

**Figure 5.**
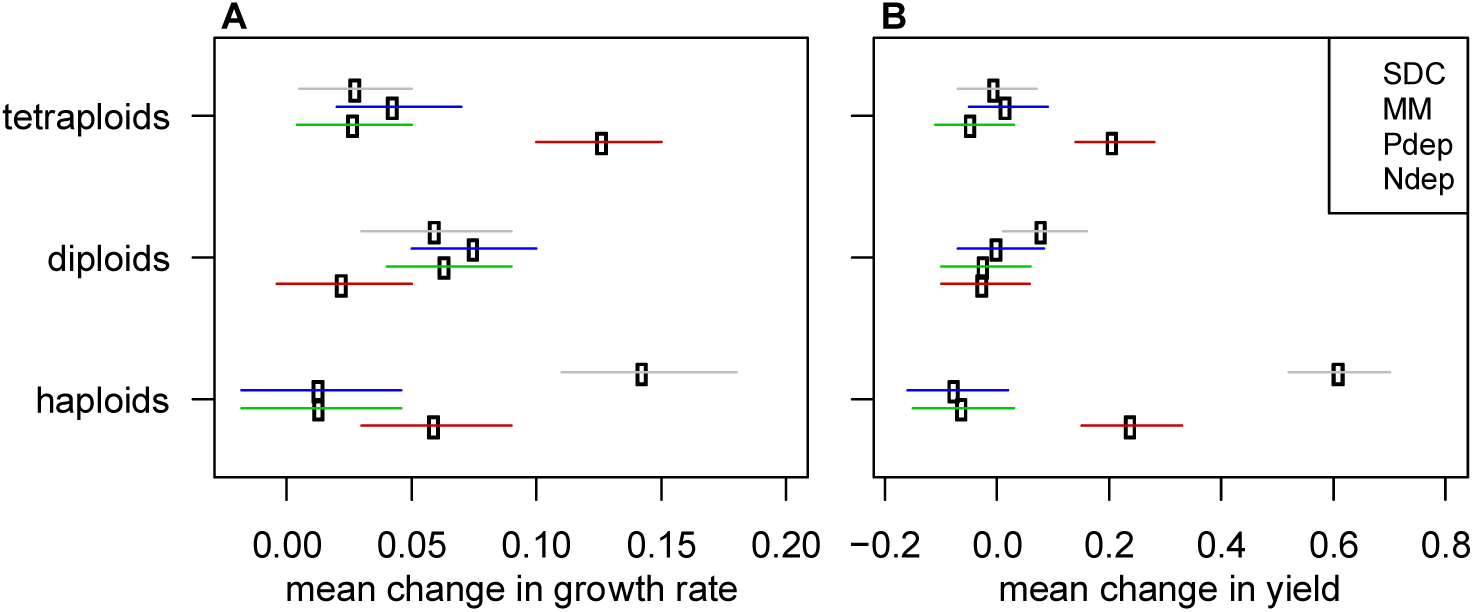
Change in fitness depended on the evolutionary environment. The mean change in A) growth rate and B) yield determined from the predicted marginal means (+/− 95% confidence intervals) in a linear-mixed effects model that treats strain background as a random effect.

Despite finding significant changes in both genome size and fitness, the observed changes in genome size across all lines did not predict the improvement in fitness when strain background and growth environment were taken into account (growth rate; haploids: F_1,78_= 2.94, *p* = 0.09, diploids: F_1,116_ = 0.92, *p* = 0.34, polyploids: F_1,155_ = 0.22, *p* = 0.64; yield: haploids: F_1,78_= 1.27, *p* = 0.26, diploids: F_1,115_ = 0.06, *p* = 0.81, polyploids: F_1,38_ = 0.34, *p* = 0.56; Figure 6). Taken together, changes in fitness did not explain the significant observed differences in the rate of ploidy drive among-strains and across environments observed throughout the evolution experiments.

**Figure 6.**
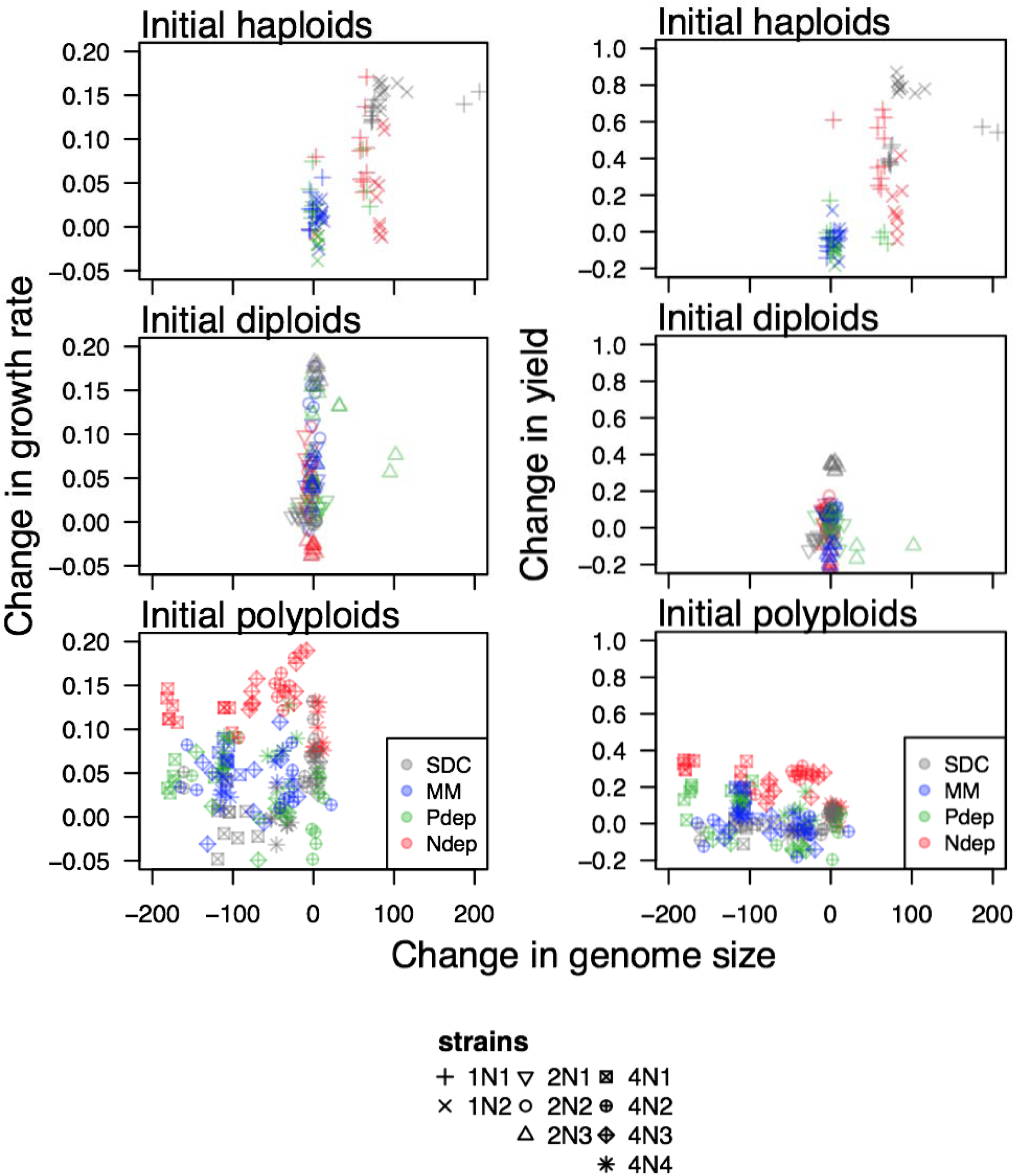
Changes in fitness generally do not correlate with the observed changes in genome size. The observed changes in genome size (negative values represent genome size reductions and positive values represent genome size gains, measured in fluorescence units) for initially haploid (top), initially diploid (middle), and initially polyploid (bottom) strains do not correlate with the observed changes in growth rate (left panel) or yield (right panel).

## Discussion

The evolutionary environment, initial ploidy, and genetic background all significantly influenced the rate of genome size change in *C. albicans*. Consistent with previous experiments in both *C. albicans* (Hickman et al. 2013; 2015) and other diverse fungal microbes (Gerstein et al. 2006; Gresham 2006; Schoustra et al. 2007; Seervai et al. 2013; Voordeckers et al. 2015), we found that the genome size changes observed over ~140 generations of batch culture evolution were nearly always towards the baseline ploidy (diploidy for *C. albicans*), a phenomenon that we term ‘ploidy drive’. By tracking genome size changes during evolution under nutrient limitation we tested whether the predicted costs of higher genome sizes (e.g., the extra P and N demands for DNA/RNA and proteins) can counteract ploidy drive. We found that the evolutionary environment did significantly influence the frequency of genome size change (Figure 2): lower ploidy levels were selected or maintained in in phosphorus-depletion (Pdep) and minimal medium (MM) while higher genome sizes were generally selected or maintained in complete medium (SDC) and nitrogen-depletion (Ndep).

We thus found contrasting results between Pdep and Ndep, inconsistent with a singular response to nutrient limitation. Rather, these results are consistent with phosphorus deprivation exerting a larger selective force counteracting ploidy drive compared to nitrogen deprivation in haploids. The ecological cost of increasing genome size under Pdep could be higher than under Ndep, so that selection against higher genome sizes would be greater in Pdep. It is also possible that other stressors that were present as the indirect effects of batch culture evolution in 96 well plates differentially influence strains evolving under Pdep and Ndep (e.g., oxygen limitation due to physical constraints limiting aeration throughout the culture blocks).

Carbon availability also cycles during batch culture evolution and it is worth considering whether this could have influenced our results, particularly as diploidization was recently been shown to be a frequent adaptive mutation under glucose limitation in *S. cerevisiae* (Venkataram et al. 2016). Our experiments used 2% dextrose for all environments (the standard amount used to culture yeast in rich/complete medium in the lab since at least 1951, Kilkenny & Hinshelwood 1951), though we have not explicitly tested whether this amount of carbon imposes a selective pressure. Preliminary experiments in *C. albicans* under 2% and 0.2% dextrose indicate that maximum growth rates are generally consistent between these two environments, but that strains reach saturation faster and at a lower population size when carbon is reduced (not shown). In the context of our experiments, this would be manifest as more time at saturation before transfer into fresh medium. Strains grown in Ndep have much slower growth rates than in the other environments (Figure 3), and thus carbon limitation (i.e., time to saturation) by this logic, would influence Ndep strains less than Pdep. However, the pattern of genome size change in Ndep was similar to strains grown in SDC, the environment where growth rates are the fastest, hence if carbon limitation is present, it is likely not causing the ploidy drive patterns observed.

Selection for lower genome sizes under Pdep (and MM) is consistent with the growth rate hypothesis, which posits that because nucleic acids contain ~9% phosphorus per unit dry mass and constitute a large fraction of organismal dry mass, there may be an increased sensitivity under phosphorus-limited conditions for higher ploidy levels (Elser et al. 1996; Sterner et al. 2002; Elser et al. 2003; Neiman et al. 2013a). Support for this hypothesis has also been found in freshwater snails, which naturally differ in ploidy level (Neiman et al. 2013b), as well as in zooplankton (Jeyasingh et al. 2015).

Genome size variant cells might arise more frequently under some conditions compared to others. The transition from haploidy to diploidy could occur more frequently under Ndep compared to MM or Pdep, so that there are simply more diploids at each mitosis and these diploids are more likely to (by chance) acquire a beneficial mutation than haploids. Similarly, the rate of genome size reduction from polyploidy towards diploidy may be less frequent under Ndep compared to MM or Pdep. Ploidy-variant causing mutations have been previously shown to be more frequent in some environments compared to others (Harrison et al. 2014). Future studies that track single-cell dynamics will help distinguish between the contributions of mutation, selection, or both on the difference in ploidy drive among environments.

Although the majority of strain backgrounds behaved quite similarly across the environments, the homozygous diploid strain (2N1) and one polyploid strain (4N1) emerged as outliers. Strain background has previously been shown to influence growth under nutrient depletion in both yeast (Magasanik and Kaiser 2002; Zörgö et al. 2013) and snails (Neiman et al. 2013b), however, the influence of genetic background on genome size stability had not been explicitly tested. Of the initially diploid strains, only lines from the homozygous diploid evolved in Pdep exhibited changes in genome size (in all cases an increase in genome size, Figure S1). Interestingly, the evolved lines that increased in genome size had reduced growth rates relative to the evolved lines that stayed diploid (mean growth of the five measured lines that increased in ploidy: 0.26 +/− 0.02; mean growth rate of the five lines that remained diploid: 0.31 +/− 0.01), a further indication that genome size transitions occur independently of increases in growth rates.

Of the initially polyploid strains, all evolved lines from the clinical strain 4N1 showed genome size reduction (Figure 2 and S1), while many evolved lines from the other clinical strain (4N2) and the two laboratory strains (4N3, 4N4) retained sub-populations of cells with their initial genome size (Figure S1). 4N1 has a unique karyotype that contains both tetrasomies and trisomies as well as two copies of isochromosome 5L (Table 1, Abbey et al. 2014). Differences in the rate of ploidy drive may be due to differences in the specific aneuploid chromosomes in 4N1 (i.e., compared to 4N3, also an aneuploid strain). Alternatively, allelic variation in a gene (or genes) involved in mutation repair or cell cycle fidelity may contribute to the rate of ploidy drive. To this end, recent analysis of heterozygous diploids revealed many DNA repair and cell-cycle regulatory genes that harbour one or more SNPs leading to allele-specific expression differences, including *POL3, RAD9, RRD1, CCN1, HAT1* and *RAD32* (Muzzey et al. 2014). Future studies are needed to shed light on particular genomic features important in promoting (or counteracting) ploidy drive.

Initial fitness was more influenced by genetic background than initial ploidy in all environments. Genome-wide homozygosity is known to reduce competitive fitness in *C. albicans* (Hickman et al. 2013), and clinical homozygous strains have yet to be obtained.

Accordingly, the three homozygous strains (1N1, 1N2 and 2N1) were consistently among the least fit strains regardless of environment (Figure 3), with the caveat that these three strains are nearly isogenic. While the nutrient limitation hypothesis predicts that higher ploidy cells will grow slower in nutrient limitation relative to lower ploidy cells, *C. albicans* polyploids typically had similar fitness as the heterozygous diploids (Figure 3), despite having larger cell size and volume (Hickman et al. 2013). Furthermore, if initial fitness dictated the rate of ploidy drive we would expect to see higher rates of ploidy drive in strains that were initially less fit. This prediction was not borne out; strain 4N1 had an increased rate of ploidy drive yet similar fitness to the other polyploid lines (with the exception of in MM). Thus the frequency of ploidy drive of polyploid lines cannot be due to differences in initial fitness.

While the majority of lines increased in fitness in after evolution (Figure 5 & S2), neither initial ploidy nor final genome size correlated with the observed changes (Figure 6). An *a priori* explanation for ploidy drive is that cells at their baseline ploidy have higher fitness than ploidy-variant cells (so that once such cells are present within a ploidy-variant population they rise to high frequency). Our results are consistent with prior empirical experiments that compared haploid and diploid growth rates in *S. cerevisiae* under nutrient limitation, where no consistent growth advantage to haploid strains has been identified (Adams and Hansche 1974; Glazunov et al. 1989; Mable 2001; Zörgö et al. 2013). Indeed, a clear selective advantage to diploidy over haploidy *per se* has only been identified in a single experiment in *S. cerevisiae*, where strains were evolved under carbon limitation (Venkataram et al. 2016). Notably, in our experiments genome size stability often ran counter to changes in fitness: haploid lines stayed haploid in MM and Pdep yet improved in fitness rate the least, the diploid lines that changed in genome size in Pdep were less fit than diploid lines that stayed diploid, polyploid lines were most stable in Ndep yet also improved in fitness the most. The force driving genome size dynamics is thus independent of the measured fitness parameters in these microbial taxa, which have streamlined genomes and low GC content, and are thus already optimized to require minimal nutrient resources for replication (Giovannoni et al. 2014, and references within).

Why ploidy drive is such a frequent and potent force across broad taxa and environmental conditions remains a mystery. As a final perplexing observation, short-term fitness patterns do not always predict long-term ploidy change. For example, *S. cerevisiae* haploid strains adapted faster than diploids in both rich medium and salt-stressed medium (Gerstein et al. 2010), yet all haploid strains converged to diploidy within 1800 generations of batch culture (Gerstein et al. 2006). We predict that had our experiments continued for longer, all of the lines would have converged on diploidy. The overarching picture is that while both the growth environment and genetic background play a role in the rate of ploidy drive, the major factor(s) that drive populations towards baseline ploidy in diverse fungal microbial taxa remains an evolutionary puzzle to be solved.

## Acknowledgements

We thank the organizers of the Elements, Genomes and Ecosystems Royal Society Theo Murphy Meeting for the initial motivation for this project, Mark McClellan and Rachel Urbitas for laboratory assistance, Jasmine Ono and Levi Morran for helpful comments, and the University of Minnesota Flow Cytometry Core Facility. The quality of the manuscript was greatly improved after full peer-review at Axios Review (Vancouver, Canada). This work was supported by R01AI0624273 grant and an ERC Advanced Award 340087/RAPLODAPT to JB. ACG was supported by a postdoctoral fellowship from the National Sciences and Engineering Research Council of Canada and a Banting Postdoctoral Fellowship from the Canadian Institutes of Health Research.

## Supporting Tables

Table S1. Tukey test results following ANOVA tests to examine the influence of environment and strain on the rate of genome size change.

**Table S2. Least square means pairwise posthoc tests to determine the influence of evolutionary environment on change in growth rate.** Degree of freedom was 342 in all tests.

**Table S3. Least square means pairwise posthoc tests to determine the influence of evolutionary environment on change in yield.** Degree of freedom was 342 in all tests.

## Supporting Figures

**Figure S1. Strain background and environment influence genome stability.** Twelve lines of each ancestral strain were evolved in complete (SDC, black points), minimal (MM, blue), phosphorus-deprivation (Pdep, green) and nitrogen-deprivation (Ndep, red) medium. Filled points represent the major peak and hollow points indicate the minor peaks in FITC intensity (see Figure 1) after ~140 generations of evolution. Top row = initially haploid lines; middle row = initially diploid lines; bottom row = initially polyploid lines. Grey boxes indicate the genome size range measured from twelve ancestral replicates for each strain.

**Figure S2. The majority of evolved lines increased in fitness.** Each point is the evolved growth rate (top panel) and yield (bottom panel) of one of 10 replicate lines evolved for each ancestral strain. Black bars indicate the median of evolved growth rates and yield. Grey bars indicate the mean of ancestral growth rates and yield as presented in Figure 3.

